# Interval-zone free-flow electrophoresis as a charge-specific dimension for native isolation of thylakoid membrane protein complexes

**DOI:** 10.64898/2026.08.27.747539

**Authors:** Lutz Andreas Eichacker, Gerhard Weber

## Abstract

Native polyacrylamide gel electrophoresis is the standard first analytical step after detergent solubilization of thylakoid membranes. It separates photosystem supercomplexes by size and shape, but it forces a polydisperse mixture of protein–detergent particles through a gel of limited pore diameter. Interval-zone free-flow electrophoresis (iZE-FFE) performs that first electrophoretic step in free solution, without ampholytes or immobilines, and returns the complexes as a liquid fraction series ordered by net charge. We describe the operational cycle of iZE, show that bromophenol blue reports the charge density of β-dodecylmaltoside micelles, and apply iZE to *Arabidopsis thaliana* thylakoid extracts from two sequential-solubilization campaigns. After a two-step digitonin then digitonin/β-dodecylmaltoside extraction (experiment 6874), SDS-PAGE of the iZE fractions is crowded: high-molecular-mass protein–lipid assemblies of photosystem I and photosystem II release the same subunits into many wells. Native PAGE of the same fractions resolves ATP synthase from PSI, a free LHCII pool, anodic PSII, and cathodic PSI, including PSI with and without LHCII. A three-step extraction with a single-pH working window (experiment 7154) confirms this charge order. Lowering residual detergent in consecutive extracts still produces partly solubilized high-mass assemblies, whereas a final β-dodecylmaltoside step releases the C2S2M2/C2S2M/C2S2 PSII series and a single PSI band. iZE is therefore a charge-first, orthogonal dimension for native thylakoid biochemistry. Native PAGE, not SDS-PAGE, is the proper second dimension while solubilization remains incomplete.

## Introduction

The photosynthetic electron-transport chain of higher-plant chloroplasts is carried by three chlorophyll-binding membrane complexes — photosystem II (PSII), cytochrome *b*_6_*f*, and photosystem I (PSI) — together with ATP synthase, the NADH dehydrogenase-like complex (NDH), and a large set of light-harvesting antenna proteins. These assemblies occupy different domains of the thylakoid membrane and change composition with light quality, state transitions, and development (Järvi et al., 2011; Anderson et al., 2012). After mild solubilization, the standard separation is native PAGE, especially blue-native (BN), clear-native, or lithium dodecyl sulfate native (LN) electrophoresis (Schägger and von Jagow, 1991; Wittig et al., 2006; Järvi et al., 2011; Arnold et al., 2014; Rantala et al., 2018). The gel reports hydrodynamic size. It does not report the net charge of the protein–detergent particle, and it begins with a collision.

Every native-PAGE lane starts as a liquid. Protein complexes, free detergent micelles, and residual membrane fragments are loaded in a dense overlay and, when the field is applied, they undergo a short free electrophoretic flight toward the stacking gel. At the liquid/gel boundary, the viscosity increases sharply. Particles with high electrophoretic mobility reach that boundary first and concentrate; smaller or less charged particles catch up while the front is still forming; small charged micelles arrive before the complexes they were meant only to solubilize. The result is a mixed, crowded entry zone. Large PSII supercomplexes and smaller PSI particles occupy the same pores at the same time. Distorted bands, material stuck in the well, and loss of labile associations are familiar costs of that geometry.

Free-flow electrophoresis (FFE) was introduced as a matrix-free, continuous electrophoretic method for cells, organelles, and membranes (Hannig and Heidrich, 1974). Intact spinach chloroplasts were purified by FFE almost fifty years ago (Dubacq and Kader, 1978), and zonal FFE remains useful for plant membrane compartments (Barkla, 2018). Those applications separate *organelles or vesicles* by surface charge. They do not separate *solubilized protein complexes*. Isoelectric focusing in free flow (IEF-FFE) has been used to measure native isoelectric points of digitonin-solubilized membrane protein complexes from Arabidopsis chloroplasts and mitochondria (Behrens et al., 2013). That mode establishes a pH *gradient* with carrier ampholytes and typically includes hydroxypropyl methylcellulose (HPMC) to suppress electroendosmosis; complexes migrate until their net charge is zero.

Interval-zone FFE (iZE-FFE) is a different experiment. It is neither continuous zone electrophoresis nor IEF. In continuous ZE, the sample is applied steadily into a medium of constant pH and is deflected while it travels with the laminar flow; that mode is the classical Hannig method for cells and organelles (Hannig and Heidrich, 1974; Barkla, 2018). In IEF-FFE, a pH *gradient* is formed with carrier ampholytes, and amphoteric analytes stop when their net charge is zero (Behrens et al., 2013). In iZE, a finite sample zone is introduced as a batch; the field is switched on while the medium moves slowly; the field is switched off; and the chamber is flushed into 96 wells (Hartmann et al., 2007; Weber and Weber, 2018).

Resolution in the protocols used here does not come from a continuous ampholyte gradient. It comes from **pH steps** forming a continuous gradient of stepped pH zones. Two or more individual media that share the same acid/base pair, but differ in pH, are loaded into the laminar flow chamber from anode to cathode. The sample is loaded on the cathodic side of the working window; an anionic particle migrating toward the anode crosses each of the pH zones; its charge and therefore its mobility change at the boundary, and the band is sharpened (Weber and Weber, 2018). The analytes do not come to rest at their pI. The same interval-plus-pH-step principle holds for different acid–base pairs. To separate Arabidopsis thylakoid membrane complexes, we used hydroxyisobutyrate (HIBA) and BisTris in the presence of digitonin (Eichacker et al., 2015; Methods). We did not use HPMC or immobilines/ampholytes in the *separation media*, so the fractions remain compatible with ultrafiltration (Eichacker et al., 2015; contrast Behrens et al., 2013; Hartmann et al., 2007). The interval zone FFE of a HeLa total-protein extract was shown to keep proteins in solution under conditions where IEF-FFE precipitated them (Hartmann et al., 2007).

We have previously reported that iZE-FFE can be followed by BN-PAGE in a methods chapter (Eichacker et al., 2015). Here we establish the charge-density control of the detergent medium, the orthogonality of iZE mobility to native PAGE, and a practical consequence of sequential thylakoid solubilization that SDS-PAGE of the same fractions cannot show. Three observations structure the work. First, the electrophoretic mobility of a small anionic dye in β-dodecylmaltoside (β-DDM) depends on detergent concentration, as expected if dye and detergent form mixed micelles whose charge per particle is diluted as more uncharged detergent is added (Helenius and Simons, 1977). Second, chlorophyll-binding thylakoid complexes occupy discrete windows along the iZE axis, and that order is the reverse of their order in native PAGE: PSII and ATP synthase are anodic, PSI is cathodic, and a free LHCII pool is resolved from both photosystems. Third, because charge separation occurs in liquid, each subsequent native-PAGE lane samples a separate charge class. SDS-PAGE of the same wells is less informative, because partly solubilized high-molecular-mass assemblies of PSI and PSII release overlapping subunit sets into many fractions. Native PAGE of those fractions is therefore the proper second dimension.

Band assignments in this preprint rest on the established subunit composition of the thylakoid complexes, on marker proteins run in the same gels, and on the literature of this laboratory and of others who have characterized the same membranes (Müller and Eichacker, 1999; Granvogl et al., 2006; Reisinger and Eichacker, 2007; Plöscher et al., 2009; Järvi et al., 2011; Behrens et al., 2013; Arnold et al., 2014; Yadav et al., 2017). Mass spectrometry of the iZE fractions, including the *stn7* and *pph1* genotypes, will be reported separately. Our finding that iZE fractions of plant stroma-lamella extracts can be used for single-particle electron microscopy of PSI–LHCI associations, including a labile contact with cytochrome *b*_6_*f* (Yadav et al., 2017), provides the structural basis for the methodological claim: the product of the first dimension is still a solution.

## Results

### Interval-zone FFE operates a five-step protocol for rapid separation of charged particles

Solubilized thylakoid extracts were introduced into the FFE chamber as a zone and resolved by iZE (1600 V, 10 °C, 0.2 mm spacer; Methods) using the interval-zone operating cycle. The cycle operates in five steps: sample injection (load), alignment of the zone between the electrodes (stack), high-voltage resolution (resolve), elution with the field off, and collection in the microtiter plates (collect), followed by fraction analysis (analyze) (Figure 1; FFE Service GmbH, 2019). In the first step (1, **Load**), the sample is pumped into the chamber with the field off while the HIBA/BisTris media, prefilled in the chamber, flow at an intermediate rate (medium). In the second step (2, **Stack**), the sample is aligned between the electrode membranes while the media flow is decreased (low) and the field is switched on. With the electric field switched on, the acid/base pair establishes the basis for the third step (3, **Resolve**). Here the acid/base pairs establish and maintain the buffer pH levels. They continuously enter the laminar flow via the adjacent inlets; the buffer components counterflow and maintain the pH *steps* from the anode to the cathode. Protein–detergent particles in the sample zone accelerate according to the charge they carry, and their mobility is modulated as they pass through each pH-step zone. Crossing a step boundary sharpens the band (Weber and Weber, 2018). In the fourth step (4, **Collect**), the field is switched off and the chamber is flushed at high flow into a 96-well plate (fraction 1 = anode). During steps 1 to 3, the laminar flow is continuously sampled to waste; during step 4, the microtiter plate is inserted into the flow stream. In step 5 (**Analyze**), chlorophyll and protein absorbance were monitored in the wells, and selected fractions were concentrated by microfiltration and analyzed on native or denaturing gels.

**Figure 1.**
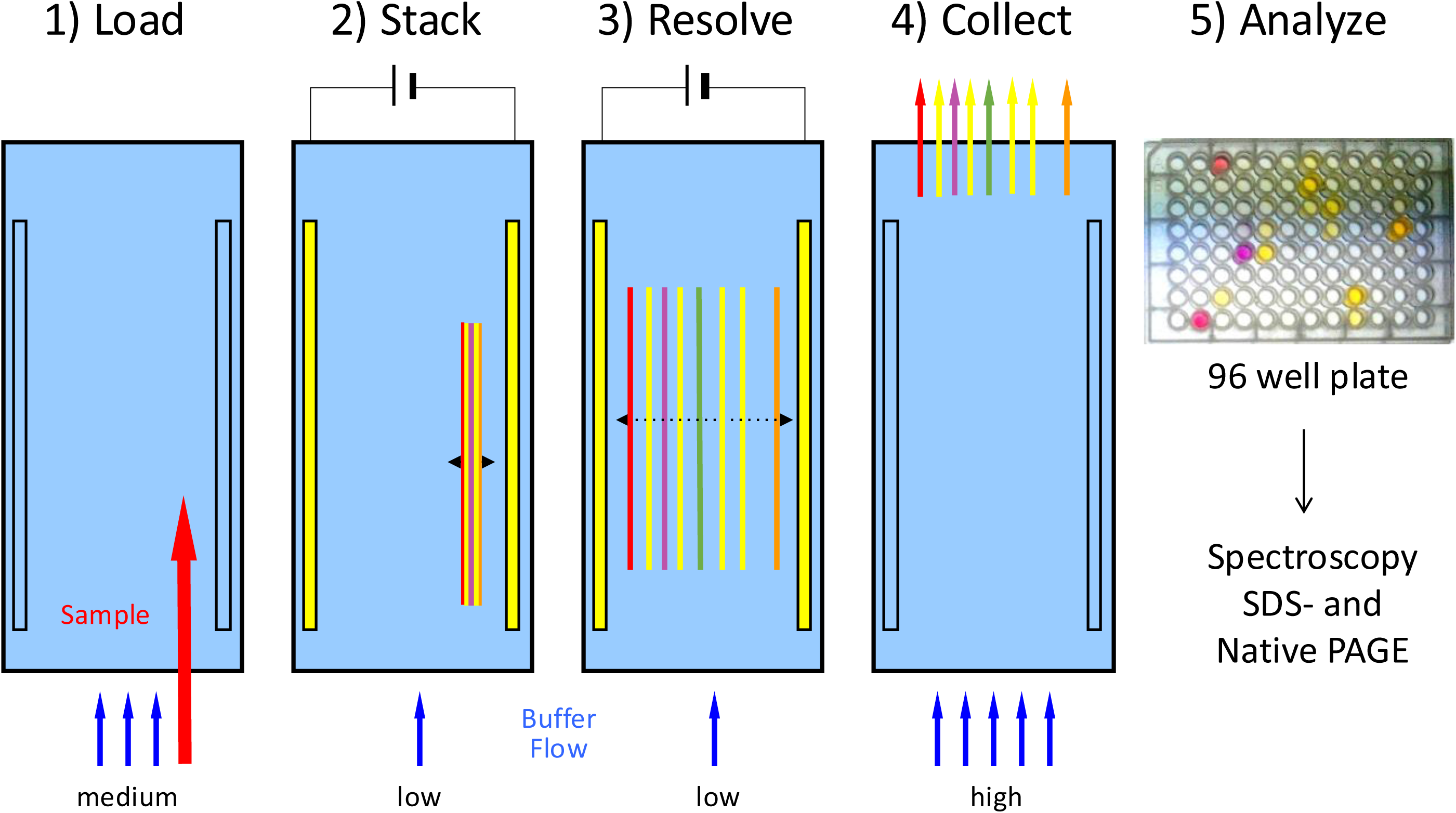
Operational principle of interval-zone electrophoresis in the FFE apparatus. A sample (red arrow) is injected by flushing a defined volume into the separation cell with a tube pump. In parallel, the FFE media flow from the bottom into the chamber at a medium flow rate (three blue arrows). The electric field is off (**1, Load**). Once the sample volume has transferred into the space between the electrodes (yellow bars), the buffer flow decreases, and the field is switched on (**2, Stack**). The free-flowing media establish pH *steps* from low/acidic (anode) to high/basic (cathode). Protein complexes loaded on the basic side of the working window accelerate toward the electrodes; complex mobility at each pH step depends on the charge state at that pH (**3, Resolve**). After the separation is established, the field is switched off, and the chamber is flushed at high flow into the 96 wells of a microtiter plate (**4, Collect**). The sample volume recovered is about 200–300 µl per fraction. The fraction contents are analyzed by spectroscopy, or the composition of proteins and protein complexes by SDS-PAGE and native PAGE (**5, Analyze**).

Under the native iZE conditions used here, chlorophyll-binding protein complexes occupied roughly 30 adjacent wells, typically within fractions 32–64, and we collected about 200–300 µl from each fraction. Complexes larger than ∼100 kDa were concentrated about eight-fold by spin filtration in ∼6 min before native PAGE. In the LN-PAGE separation typically used for experiment 7154, no Coomassie addition was required to charge the protein complexes during the run, and the polyacrylamide gel did not require destaining or electroelution before subsequent measurement.

### Bromophenol blue reports the charge density of DDM micelles

Bromophenol blue (BPB) is a small, strongly anionic sulfonephthalein. It is **not** an amphoteric pI marker. In the pH range of the FFE chamber (pH ∼3–9), the sulfonate remains negatively charged. Even at pH 3.0–4.6, the color transition reflects phenolic ionization, not loss of electrophoretic charge. A constant amount of BPB was dissolved in β-DDM and subjected to the same iZE program at 0.25, 0.5, 1, 2, 4, 6, 8, and 10 mM DDM (Figure 2). Anodic mobility of BPB fell as DDM concentration increased. All DDM concentrations are above the critical micelle concentration of DDM in water, 0.17 mM (0.0087%) (VanAken et al., 1986). In the titration, we determined the inflection point for BPB mobility by differentiation at about 2.2– 2.5 mM DDM. Considering an N_agg_ of 100–140 for DDM, the formula [micelle] = ([DDM] − CMC)/N_agg_ can be applied. The inflection point reflects the effective concentration for BPB retardation by DDM micelles, not the thermodynamic CMC of pure DDM.

**Figure 2.**
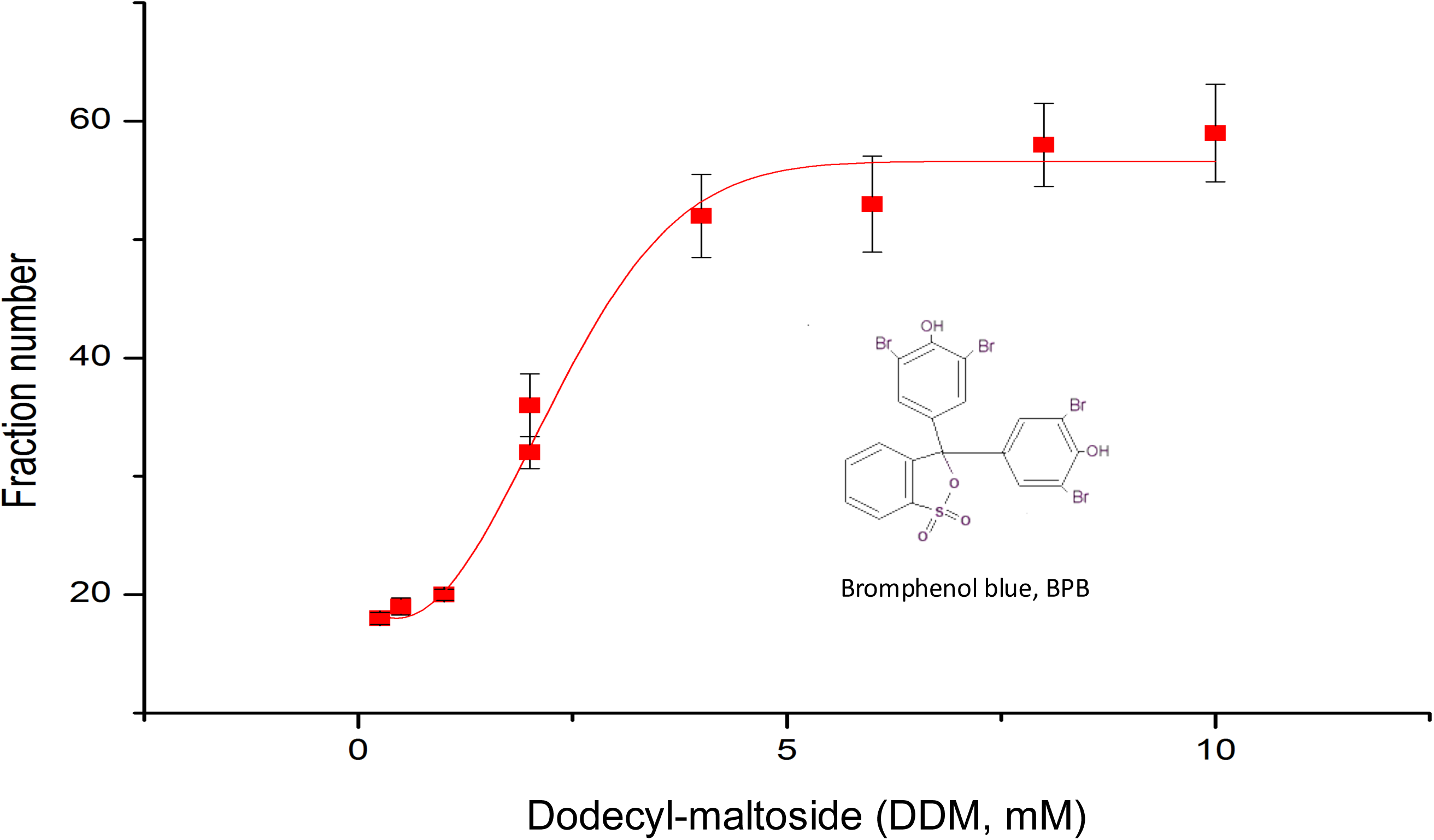
Charge density separation of β-dodecylmaltoside micelles reported by bromophenol blue. Detergent vesicles of increasing DDM concentration (0.25, 0.5, 1, 2, 4, 6, 8, and 10 mM) were labeled with a constant concentration of bromophenol blue. Samples were separated by iZE and collected as 300 µl fractions (wells 10–70). The absorbance of BPB distribution per fraction (mean ± SD of three technical replicates) is plotted against DDM concentration. Anodic mobility decreases as DDM concentration rises.

The same iZE runs contained the coloured pI-marker mix used to read the pH steps (Šlais and Friedl, 1994; Methods). Absorbance at 420, 515, and 595 nm showed that the amphoteric dyes stayed in place, while only BPB was retarded. In two further controls, increasing digitonin concentrations produced **no** BPB shift, indicating that the micellar interiors of DDM and digitonin are not equivalent solvents for the dye. Orange G, an always-anionic azo dye, produced **no** DDM-dependent shift. BPB peak intensity did not increase with DDM, so the dye is not aggregating; the extra hydrodynamic mass comes from the detergent.

The observation is the free-solution analog of charge-shift electrophoresis (Helenius and Simons, 1977). BPB partitions into DDM micelles. Additional uncharged detergent increases the number of detergent molecules per charged dye (or per charged protein–detergent particle), lowering the charge-to-friction ratio. Two practical consequences follow. First, the detergent concentration in the FFE medium is not a spectator variable; it sets the mobility of every micellar particle in the run. Second, after solubilization, the extract contains a large population of protein-free, lipid- and detergent-rich micelles. Those particles are small and charged. In a gel, they reach the polyacrylamide front before the photosystem complexes. In iZE, they occupy their own charge window and can be left out of the lanes loaded later.

Why only BPB, of the dyes in the mix, reports DDM is experimental rather than fully structural. The Šlais markers are substituted aminomethylphenols (and, in the acidic range, azo dyes) designed as hydrophilic ampholytes with low hydrophobicity so that they focus at a defined pI (Šlais and Friedl, 1994; Štastná et al., 2005). BPB is a halogenated sulfonephthalein that remains anionic throughout the run and partitions into DDM. Digitonin micelles apparently do not take it up. We do not claim a binding site.

### The pI-marker set locates the pH steps of the iZE medium

Before loading protein complexes, the colored pI-marker mix is separated under the same iZE program, and the pH of the collection plate is measured (Figure 3). The markers occupy discrete wells. The pH recorded after the separation reports the steps that the HIBA/BisTris media actually establish in the chamber: an anodic acidic zone, a plateau near pH 5.5 (approximately fractions 17–41), a plateau near pH 6.2 (approximately fractions 43–64), and a cathodic plateau near pH 7.0. This is the operational signature of interval-zone electrophoresis in pH steps (Weber and Weber, 2018). The markers are used to locate those steps. They are not used to assign isoelectric points to the protein–detergent particles that follow.

**Figure 3.**
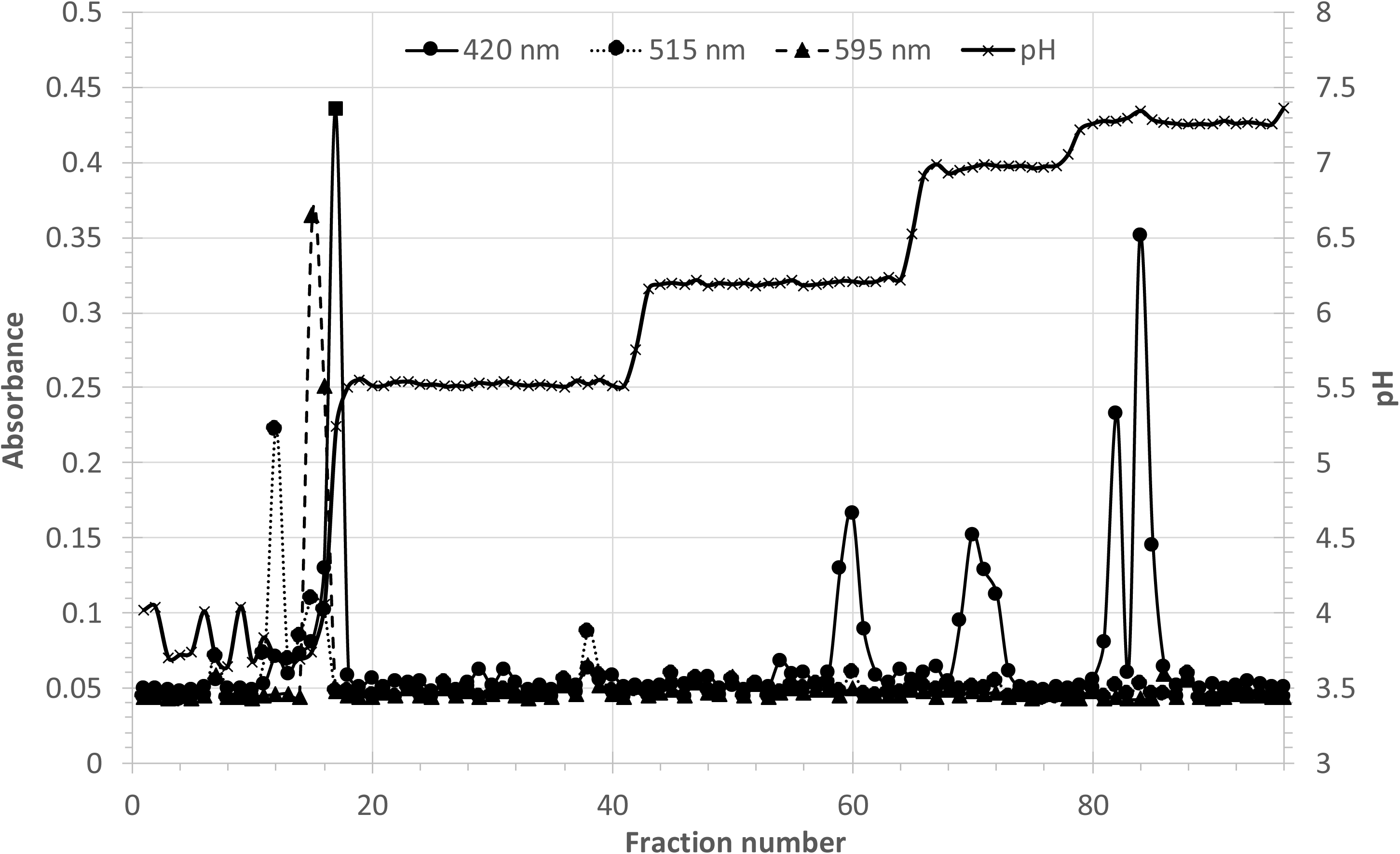
Separation of the pI-marker set and pH steps of the iZE medium. A coloured pI-marker mix (pH 4.0, 4.8, 5.3, 6.4, 7.5, 8.5 and 10.1 plus SPADNS; Šlais and Friedl, 1994) was separated under the same iZE program used for the protein extracts. Absorbance was recorded at 420 (4.0, 5.3), 515 (4.8) and 595 nm (6.4, 7.5, 8.5, 10.1) to locate the marker bands relative to the pH steps. The pH was measured in the collection plate after iZE separation. The media establish steps near pH 5.5 (approximately fractions 17–41), pH 6.2 (approximately fractions 43–64), and pH 7.0. The pI markers report mobility toward the next anodic pH step; they are not used to assign pI values to the protein complexes.

### Separation of thylakoid membrane protein complexes upon two-step solubilization in the classical multi-step pH window

In the initial phase of the study, we followed the experiment labeled #6874 and the solubilization described in Eichacker et al. (2015), using the classical iZE concept of sufficient pH steps to focus and separate the complexes (Weber and Weber, 2018). Arabidopsis thylakoids were extracted first in 16 mM digitonin. The centrifugal supernatant of that step (solubilization 1) contains the complexes released from the more accessible membrane domains. The membrane pellet recovered after that centrifugation was then extracted in a mixture of 8 mM digitonin plus 8 mM β-DDM (solubilization 2). We separated each supernatant by iZE in HIBA/BisTris media containing 0.1% (w/v) digitonin (Figure 4 and Supplemental Figure 1).

**Figure 4.**
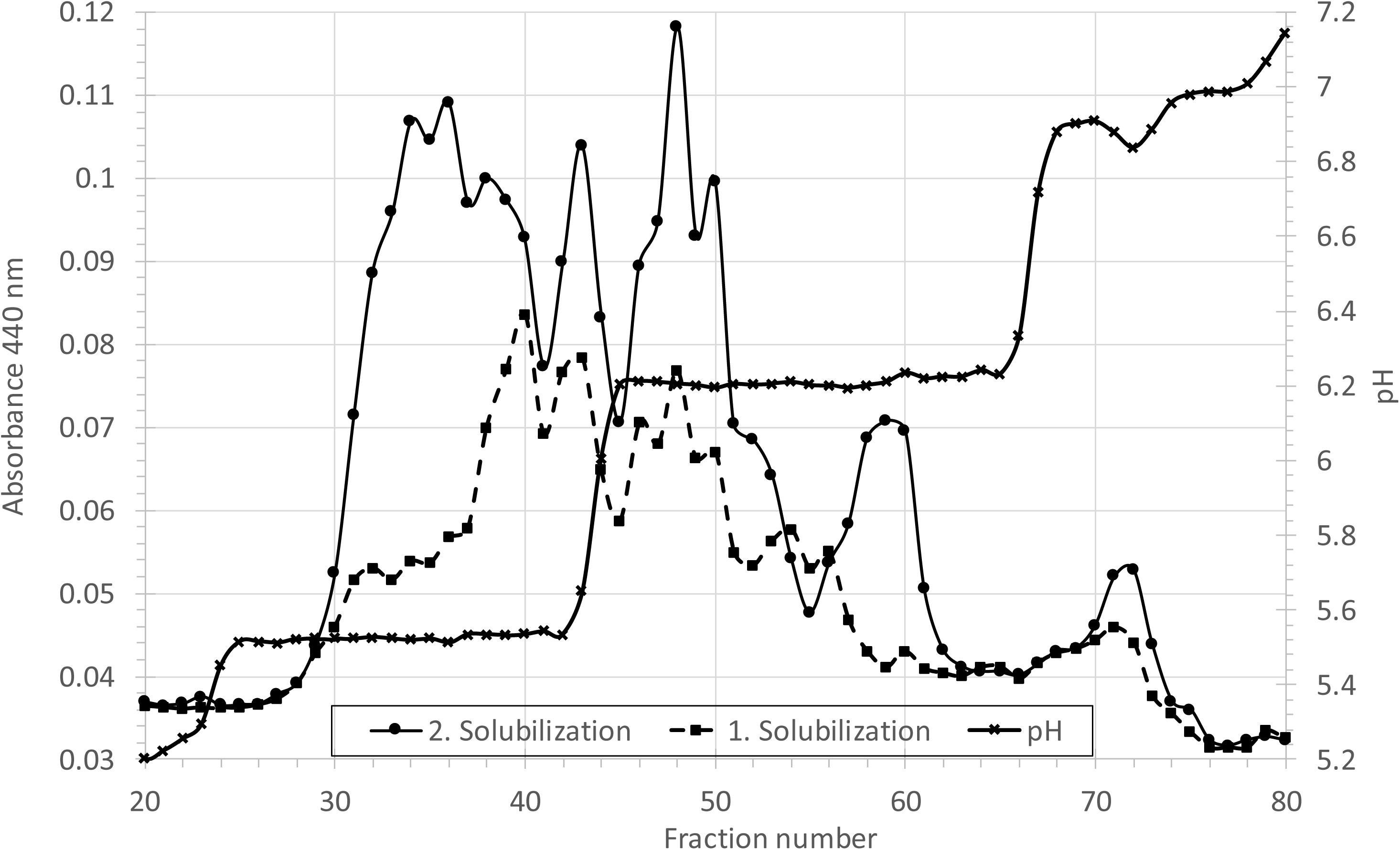
Absorbance of Chl-protein complexes and pH steps after iZE from two-step solubilization extracts of thylakoid membranes. Arabidopsis thylakoids were solubilized in two consecutive steps as described (Eichacker et al., 2015). **Solubilization 1:** centrifugal supernatant after 16 mM digitonin. **Solubilization 2:** the membrane pellet recovered after step 1, extracted in 8 mM digitonin plus 8 mM β-DDM. Each supernatant was separated by iZE (0.1% digitonin in HIBA/BisTris media, 1600 V, 4.5 min). Absorbance at 440 nm indicates the location of the chlorophyll-binding photosystem complexes. The pH was measured in the microtiter collection plate after iZE. The pH steps in the separation window of the membrane protein complexes are at 5.5, 6.2, and about 6.85.

The absorbance at 440 nm was recorded in the consecutive fractions as a direct measure of the chlorophyll-binding particles present in each extract (Figure 4). The two traces are structured rather than a single broad peak, and they are detergent-specific, as expected if digitonin and the digitonin/DDM mixture release different mixtures of complexes (thylakoid stroma-lamella-enriched versus grana-enriched; Järvi et al., 2011). The pH measured in the collection plate *after* the iZE of the protein complexes still shows the 5.5 and 6.2 steps in the chlorophyll window (Figure 4). The complete pH distribution in all 96 wells is disclosed (Supplemental Figure 1). The pH distribution in the separation window matches the pI-marker plate (Figure 3). Several chlorophyll protein complex maxima sit at the edges of those steps, which is the operational signature of band sharpening at the boundaries (Weber and Weber, 2018). Because detection is intrinsic, there is no staining delay and no need to sacrifice a gel to determine into which fraction the protein complexes were separated. The profile also identifies wells without pigmented complexes, allowing you to focus on the relevant fractions before concentration.

### SDS-PAGE analysis of the iZE fractions in experiment #6874 does not display the distribution map of the solubilized native complexes

To analyze protein subunit distribution, we denatured and separated 10 µL of the same iZE fractions by SDS-PAGE (Figure 5). Protein subunits were visualized with the SilverQuest Silver Staining Kit (Invitrogen, Thermo Fisher Scientific, catalog no. LC6070) using the microwave protocol supplied with the kit. The protein subunit labeling is necessarily rough. It is based on the known subunit composition of the complexes after SDS dissociation, on marker proteins (M) run in a separate lane, and on the unfractionated sample (S) loaded beside the iZE series (Müller and Eichacker, 1999; Granvogl et al., 2006; Reisinger and Eichacker, 2007; Plöscher et al., 2009; Arnold et al., 2014). PSI is labeled **I**, PSII **II**, ATP synthase **IV**, and the major light-harvesting protein of photosystem II **L2**.

**Figure 5.**
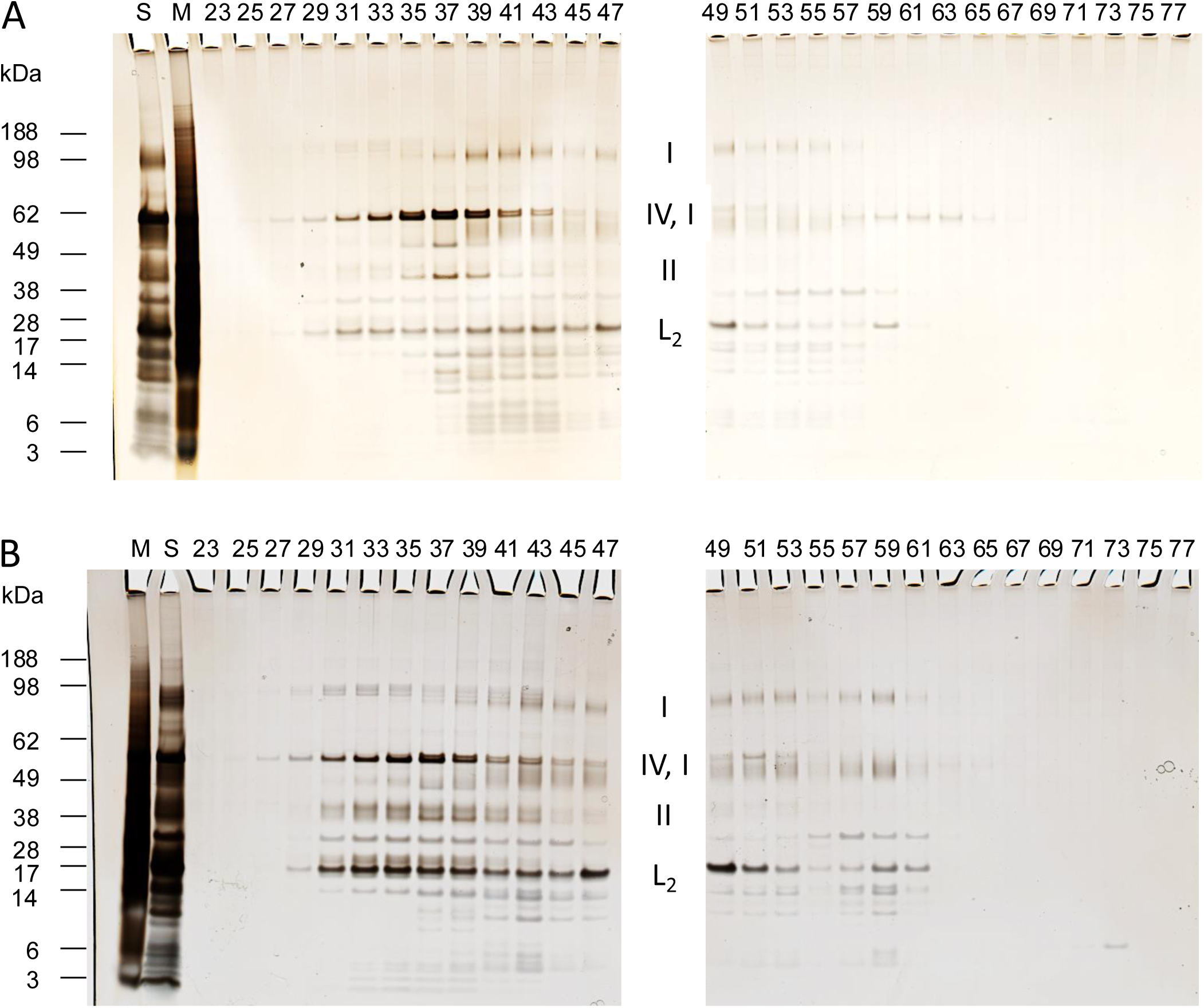
SDS-PAGE of iZE fractions from the two-step solubilization of thylakoid membranes. Protein complexes in the odd-numbered iZE fractions 23–77 from solubilization 1 (**A**) and solubilization 2 (**B**) were dissolved in SDS, separated on NuPAGE 4–12% Bis-Tris gels, and protein subunits stained with the SilverQuest Silver Staining Kit (Invitrogen, Thermo Fisher Scientific). The mass (kDa) of marker proteins **M** and of the unfractionated sample, **S**, were co-separated with the sample proteins. Protein subunit labels are assignments from the known polypeptide composition of the protein complexes and from the marker set: **I**, photosystem I; **II**, photosystem II; **IV**, ATP synthase; **L2**, LHCII. The labeling is speculative, as overlap among fractions is very high because the release of protein subunits from high-molecular-mass, partly solubilized PSI and PSII assemblies overlaps with protein subunit release from fully solubilized complexes (see Figure 6).

From the subunit composition, a native separation of different complexes into a series of consecutive fractions is difficult to resolve. There is a high degree of overlap among fractions. After 16 mM digitonin (Figure 5A), the strongest patterns are consistent with PSI and ATP synthase; after the second extraction of the residual membranes (Figure 5B), PSII and LHCII become more prominent, with L2 appearing both with PSI and as a separate pool. The detergent-specific contrast in protein subunit distribution across fractions in the two SDS gels (Fig. 5A and B) is real, but it is not a clean map from which the location of the differently charged protein complexes can be determined. The reason becomes apparent only when the same fractions are examined as intact complexes by native PAGE.

### Native PAGE of the same fractions shows that iZE effectively separated the solubilized membrane protein complexes

When we separated the same iZE fractions by blue-native PAGE, the protein complexes stained with colloidal Coomassie G-250 according to Kang et al. (2002) showed a much cleaner picture (Figure 6). We present the two BN-PAGE gels in the same order as the SDS-PAGE gels. In panel A, digitonin shows a clear preference for solubilizing photosystem I, PSI (Figure 6A, I, fractions 47-59) and ATPase holocomplex protein complexes (Figure 6A, IV, fractions 35-41). Interestingly, the distribution of the Cytochrome b6f (Cyt b6f) complex appears correlated with the PSI complex (Figure 6A, I and V). Also, the Cyt b6f band only correlates with the mobility of the more cathodic PSI complexes (Figure 6A, fractions 51 to 59), whereas no band correlation is evident with the PSI complexes binding LHCII protein (Figure 6A, fractions 47 to 52). The Digitonin solubilization also shows a smaller quantity of PSII supercomplexes (Figure 6A, II, fractions 35-39) and of light-harvesting protein in trimeric complexes (Figure 6A, L2, fractions 47 to 51). The BN-PAGE staining intensity indicates a clear concentration of ATPase in fractions 37-39, of PSI-L2 in fractions 49-51, and of PSI in fractions 55-57 (Figure 6A, IV, IL2, I). In contrast, residual membrane solubilization in a mixture of Digitonin and DDM shows and increase in the PSII and in the LHCII yield (Figure 6B, II, L2). We noted that the fraction numbers in which protein complexes of ATPase, PSII, and LHCII trimers were separated remained constant in iZE separations A, and B (Figure 6). However, the PSI complexes and Cyt b6f shifted cathodically by about five fractions to fractions 51-55 for PSI-L2, and fractions 55-61 for PSI and Cyt b6f (Figure 6B). This could indicate that the charge state of PSI is sensitive to DDM exposure or that in the presence of DDM the PSI complexes isolated more from the grana membrane section of the thylakoid membrane contain PSI in a different charge-state organization. We also noted that the iZE separation of the Cyt b6f complex followed the cathodic shift of the PSI complex, corroborating a potential interaction between both complexes as proposed earlier (Yadav et al., 2017). The BN-PAGE panel B of the two step solubilization experiment (Figure 6B) resembles the same gel previously published as Fig. 1 of Eichacker et al. (2015) and is shown here from the original scan, not from the Springer typesetting.

**Figure 6.**
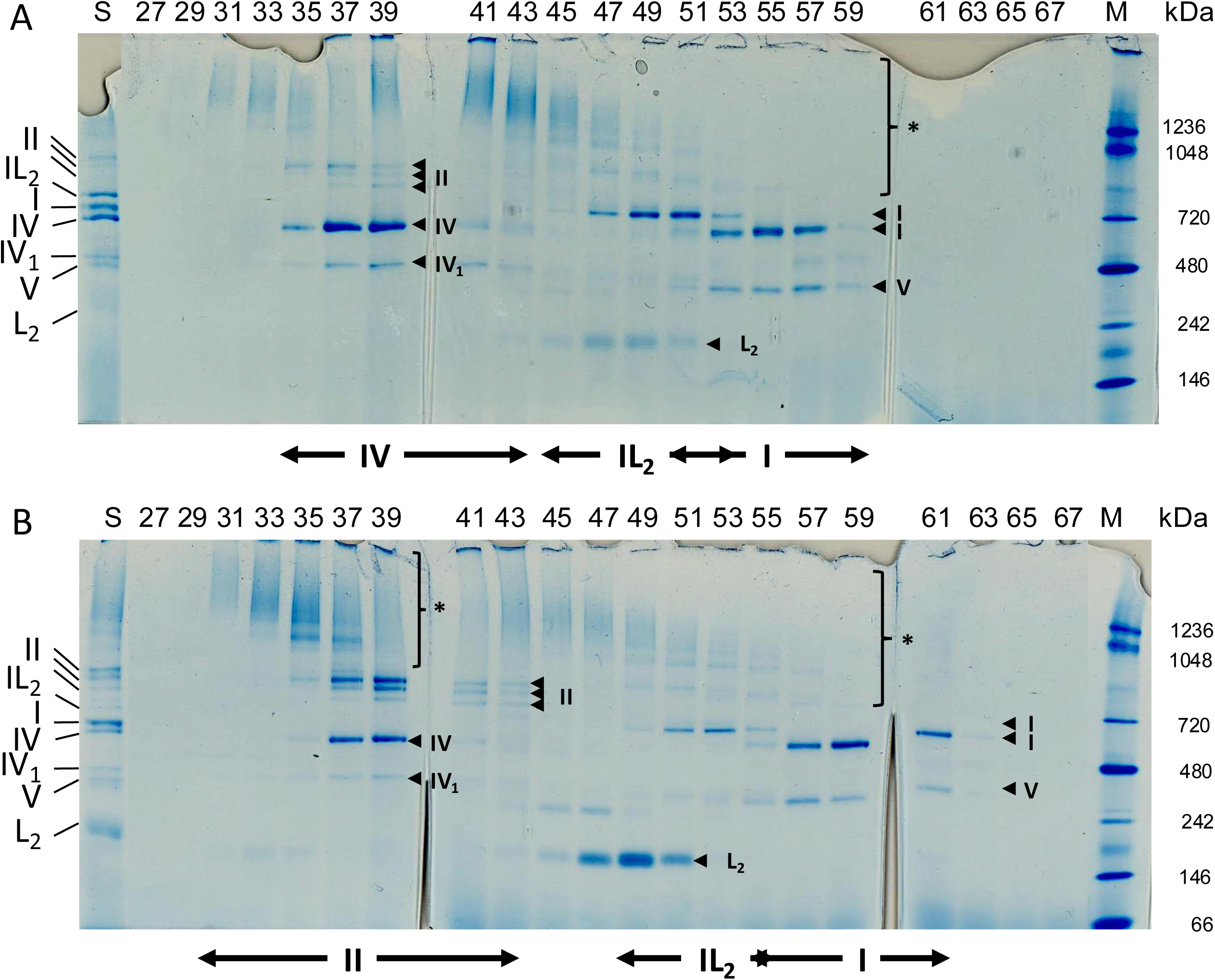
Blue-native PAGE of iZE fractions from the two-step solubilization of thylakoid membranes. Odd-numbered fractions 27–67 from solubilization 1 (16 mM Digitonin, **A**) and solubilization 2 (8 mM Digitonin plus 8 mM β-DDM, **B**) were concentrated and separated on NativePAGE 3–12% Bis-Tris gels with 0.24 mM Coomassie G-250 in the cathode. Complexes were stained with colloidal Coomassie G-250 (Kang et al., 2002). Protein complex assignments are provided from the mass spectrometry analysis (not shown) of the LN-PAGE gels as CF_0_CF_1_ ATP synthase (IV), and CF_1_ ATP synthase (IV1), PSII (II), PSI (I), cytochrome *b*_6_*f* (V), LHCII (L2), and PSI-LHCII (IL2). The asterisks mark high-molecular-mass Coomassie-stained protein complexes of PSI and PSII that increase in apparent mass toward the anode. The range of protein complex separation in iZE is indicated by arrows at the bottom of the native PAGE gels. The mobility of protein complexes from the membrane extract not separated by iZE (S), and of the NativeMark standard (kDa) (M), are provided, and complexes are labeled. **Panel B was previously published as** Fig. 1 **of Eichacker et al.** (2015) **and is reproduced here from the original scan.**

We concluded from the experiment that ATP synthase (IV) separates well from PSI (I). A separate L2 pool also becomes evident, especially in the second solubilization. PSII and ATP synthase are separated anodic in iZE; while PSI and Cyt b6f were identified with a cathodic mobility. The intensity of the protein band staining by colloidal Coomassie furthermore provides some indication of the quantitative nature of the split of PSI into a complex with and without L2 bound. The split is visible in both solubilization panels, but PSI-L2 is clearly enriched in the Digitonin-solubilized thylakoid membranes (Figure 6A versus B). Dimeric cytochrome *b*_6_*f* (V) appears in a coordinated way in the same iZE fractions as PSI. The first extraction releases more ATP synthase and PSI; the second extraction releases more PSII and L2.

We furthermore noted that the same complexes sort in **opposite** order on the two axes: charge in iZE and molecular mass or size in BN-PAGE. That is the definition of an orthogonal two-dimensional separation: charge in liquid, mass/size in the gel. It is also the opposite of the naive expectation that “more electrophoresis always means a mobility further down the gel.” In free solution, electrophoretic mobility scales with charge and inversely with friction. In a restrictive gel, size dominates gel retardation. PSII supercomplexes are highly charged *and* large; they clearly excel in the liquid but lose in the gel race. PSI does the reverse.

An additional feature of the native gels is a high degree of diffusely Coomassie-stained protein at high molecular mass. These protein structures shift upward — they increase in apparent mass — toward the anode. Both PSI and PSII populate these diffuse, high-molecular-mass forms. These clouds of protein assemblies appear to be the main reason the clean separation native PAGE produced could not be seen in SDS-PAGE, because the high-mass PSI and PSII assemblies, once denatured by SDS, released the same subunits into the gel lanes, overlaying the anodic FFE fractions of the fully solubilized protein complexes.

We interpret these clouds as a mixture of incompletely solubilized complexes as the detergent is consumed by thylakoid membrane lipids during the solubilization of the specific protein complexes. As the free detergent concentration decreases, complexes are released with different numbers of peripheral lipids and, potentially, additional proteins. All of those particles populate the centrifugal supernatant of extract 1 and, after the residual pellet is re-extracted, of extract 2. This phenomenon is more strongly detectable with digitonin alone for PSI complexes, and when DDM is added to digitonin in the second step, the phenomenon is more strongly detected for PSII complexes. This indicates that decreasing detergent activity increases the percentage of higher-molecular-weight lipid-protein complex assemblies. A related titration, in which DDM was increased on a digitonin-solubilized thylakoid membrane, and the products were examined directly by native PAGE, confirmed this analysis as it produced the same qualitative picture: a continuum of high-mass assemblies collapse toward discrete complex bands as detergent becomes sufficient (Eichacker, unpublished).

If that interpretation is correct, the clouds differ in charge because more charged lipids remain associated with the complexes, and/or because a higher number of charged proteins remain present in the more anodic particles. However, for the separation potential of iZE, our results show that iZE separates all the charged classes of protein complexes well. Native PAGE then separates them by size, while SDS-PAGE conflates them.

### Separation of the thylakoid membrane protein complexes upon a three-step solubilization in a single working-pH window by iZE

In our two-step solubilization experiment #6874, we followed the classical multi-step pH design for iZE. Native PAGE, however, typically uses one pH to set the charge state of the proteins in the gel. We further reasoned that dropping the working pH from about 6.2 toward 5.5 would also slow the mobility of the negatively charged, anodic protein complexes and protein–lipid clouds condensing the separation space. We therefore asked whether the complexes would still resolve if the iZE working window was effectively represented by a single pH. We implemented this investigation in experiment #7154. We combined the study with a three-step solubilization protocol to lower the first Digitonin extraction, lower the DDM concentration in the second solubilization step, and complete the extraction by a pure DDM solubilization step of the residual membranes. (Methods): (A) 10 mM digitonin; (B) 10 mM digitonin plus 5 mM β-DDM on the A pellet; (C) 20 mM β-DDM on the B pellet. Each extract was loaded on iZE twice. The overlay of the two independent A_440_ traces shows that the same wells can be pooled. As in the previous study, the FFE media contained 0.1% digitonin and 250 mM mannitol. The pH recorded in the collection plate after the sample runs plateaus at ∼6.4 across fractions 23–68 (Figure 7) for the separation window, while in the previous experiment “#6874 it was the 5.5 and 6.2 steps that were retained after experiment 6874 (Figure 4). We, however, also document here the complete pH profile of all 96 iZE fractions (Supplemental Figure 2), while the 440 nm traces of extracts A, B and C in Figure 7 focus on the separation window only, and act as reference for the protein-complex bands on the corresponding native gels (Figures 8–10). ATP synthase (IV), PSII (II), PSI (I), LHCII (L2) and cytochrome *b*_6_*f* (V) are assigned to the A/B/C series on the basis of those gels (Figures 8–10). Independent loads of the same extract were superimposed closely enough to pool fractions for native PAGE.

**Figure 7.**
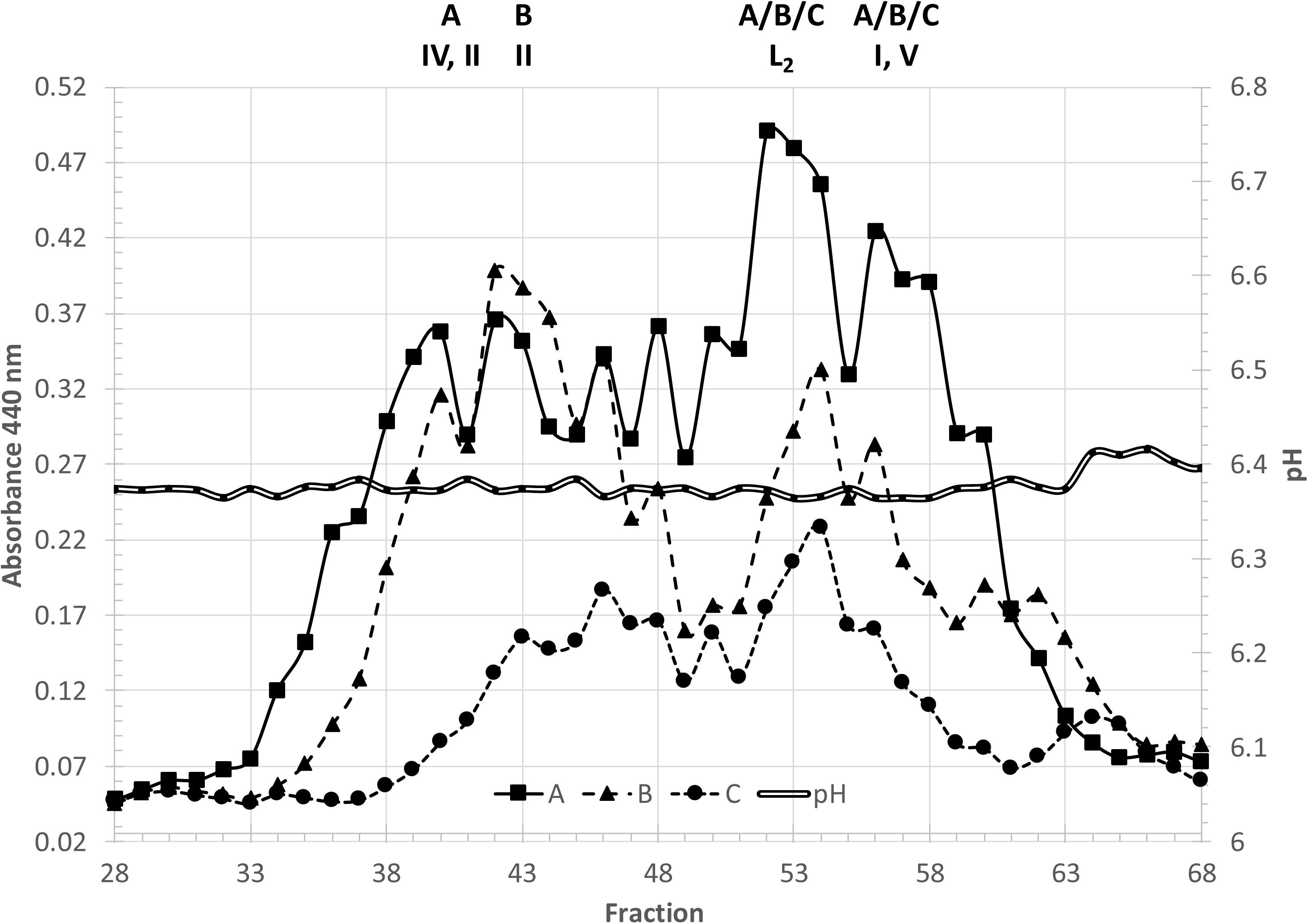
Separation of Chl-protein complexes by iZE-FFE upon three-step solubilization using Digitonin and β-DDM at pH 6.38. Arabidopsis thylakoids were extracted in three consecutive steps: 10 mM digitonin, **A**; 10 mM digitonin plus 5 mM β-DDM on the A pellet, **B**; and 20 mM β-DDM on the B pellet, **C**, and each supernatant was separated by iZE. The absorbance of Chl bound to photosystem complexes of the three extracts was recorded at 440 nm, and spectra were overlaid in the graph. The pH was measured after the sample runs and plateaued at ∼6.4 across the chlorophyll protein window (fractions 23–68). Protein complex assignments are provided from the mass spectrometry analysis (not shown) of LN-PAGE gels (Figures 8–10). ATP synthase (IV), and PSII (II) were recorded in extract A (around fraction 40/42), and of PSII (II) in extract B (around fraction 40-50); while the majority of PSI (I) and cytochrome *b*_6_*f* (V) (around fractions 56-62), and of LHCII (L2) complexes (around fractions 52/56) were recorded in all three membrane extracts A/B/C.

**Figure 8.**
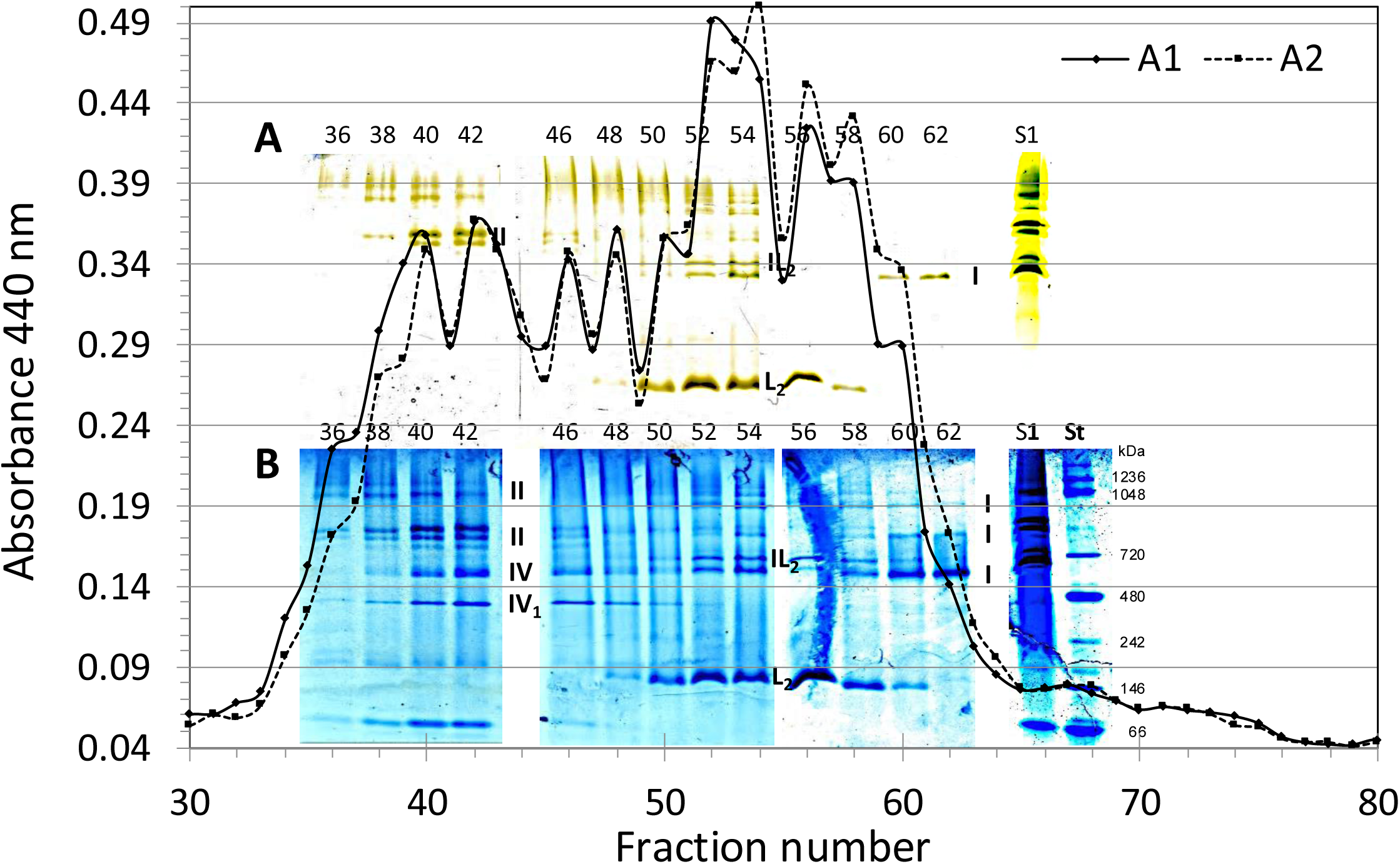
2D iZE-FFE/LN-PAGE fractionation of a 10 mM Digitonin thylakoid membrane extract. **(A).** Thylakoid membranes (400 µg Chl) were extracted in 10 mM Digitonin (methods). Two technical replicates, **A1** and **A2**, of 100 µl each were separated by iZE-FFE. The absorbance of fractions A_440_ was recorded. Even-numbered fractions 36 to 62 were concentrated by microfiltration (100k) for LN-PAGE separation (Arnold et al., 2014). The gels were white-light scanned: chlorophyll (unstained) **A**; and the same gel after colloidal Coomassie staining (**B**; Kang et al., 2002). The mobility of protein complex bands was compared to unfractionated A extract **S1** and the molecular weight NativeMark standard **St** (Methods). Protein complexes of PSI (I), PSII (II), ATPase (IV), light-harvesting protein of PSII (L2), and L2 bound to PSI (IL2) were labeled in the LN gel.

**Figure 9.**
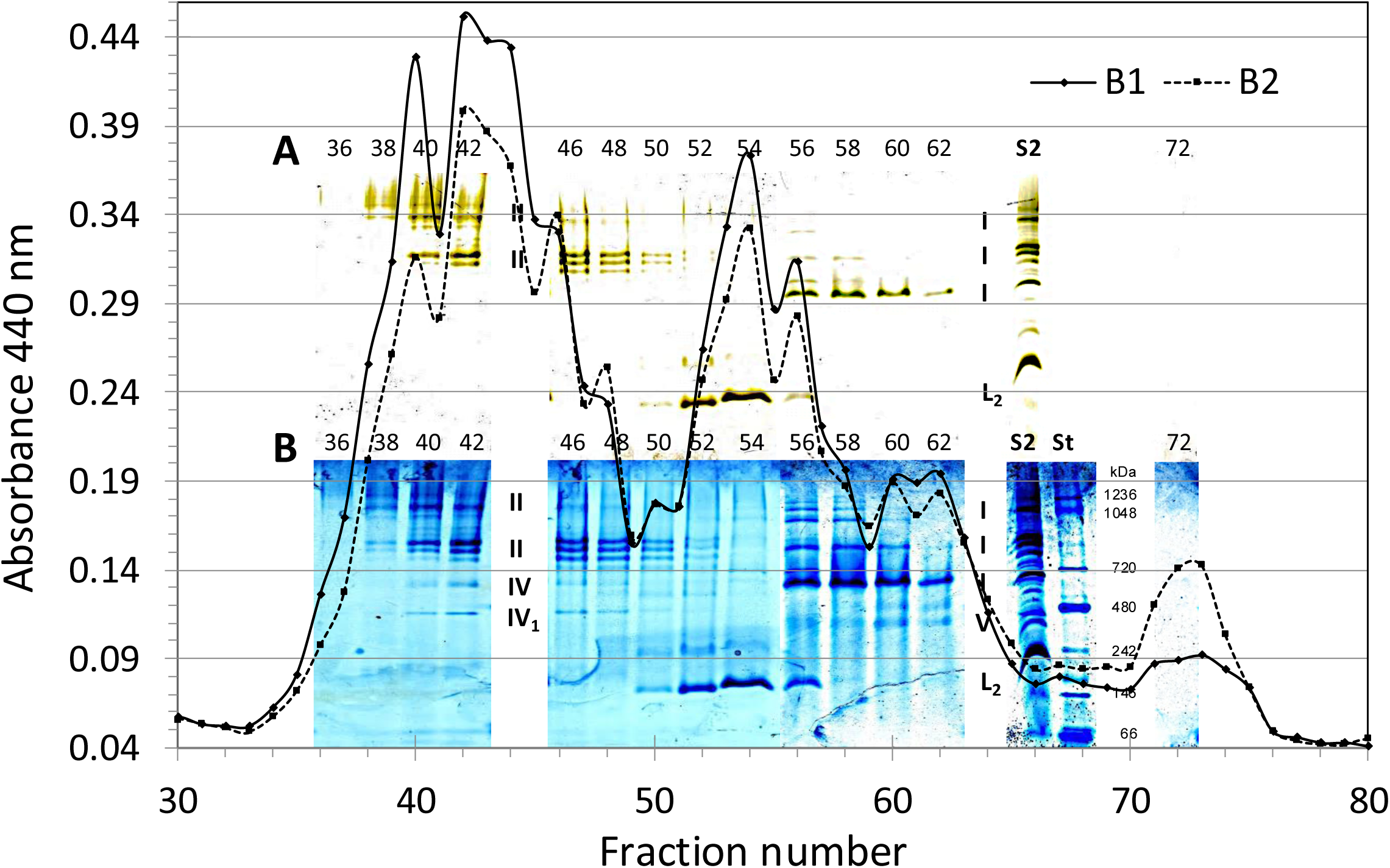
2D iZE-FFE/LN-PAGE fractionation of thylakoid membranes not extracted in step A using 10/5 mM digitonin/β-DDM (B). The thylakoid membranes not solubilized upon extraction A (Fig. 8) were recovered by centrifugation (30 min, 25 krcf, 10 °C). Membranes were extracted using 10 mM digitonin plus 5 mM β-DDM from the A pellet. Separation by iZE-FFE was conducted in two technical replicates, **B1** and **B2**. The two A_440_ overlay microplate scans from B1 and B2 were compared with LN-PAGE as described (Figure 8, Methods) and with unfractionated B extract **S2** and the molecular weight NativeMark standard **St**. Even-numbered fractions 36 to 62 were concentrated by microfiltration (100k) for LN-PAGE separation (Arnold et al., 2014). The gels were white-light scanned: chlorophyll (unstained) **A**; and the same gel after colloidal Coomassie staining (**B**; Kang et al., 2002). Protein complex band labels are assigned as shown in Fig. 8; the cytochrome *b*_6_*f* (V) complex is labeled in addition.

**Figure 10.**
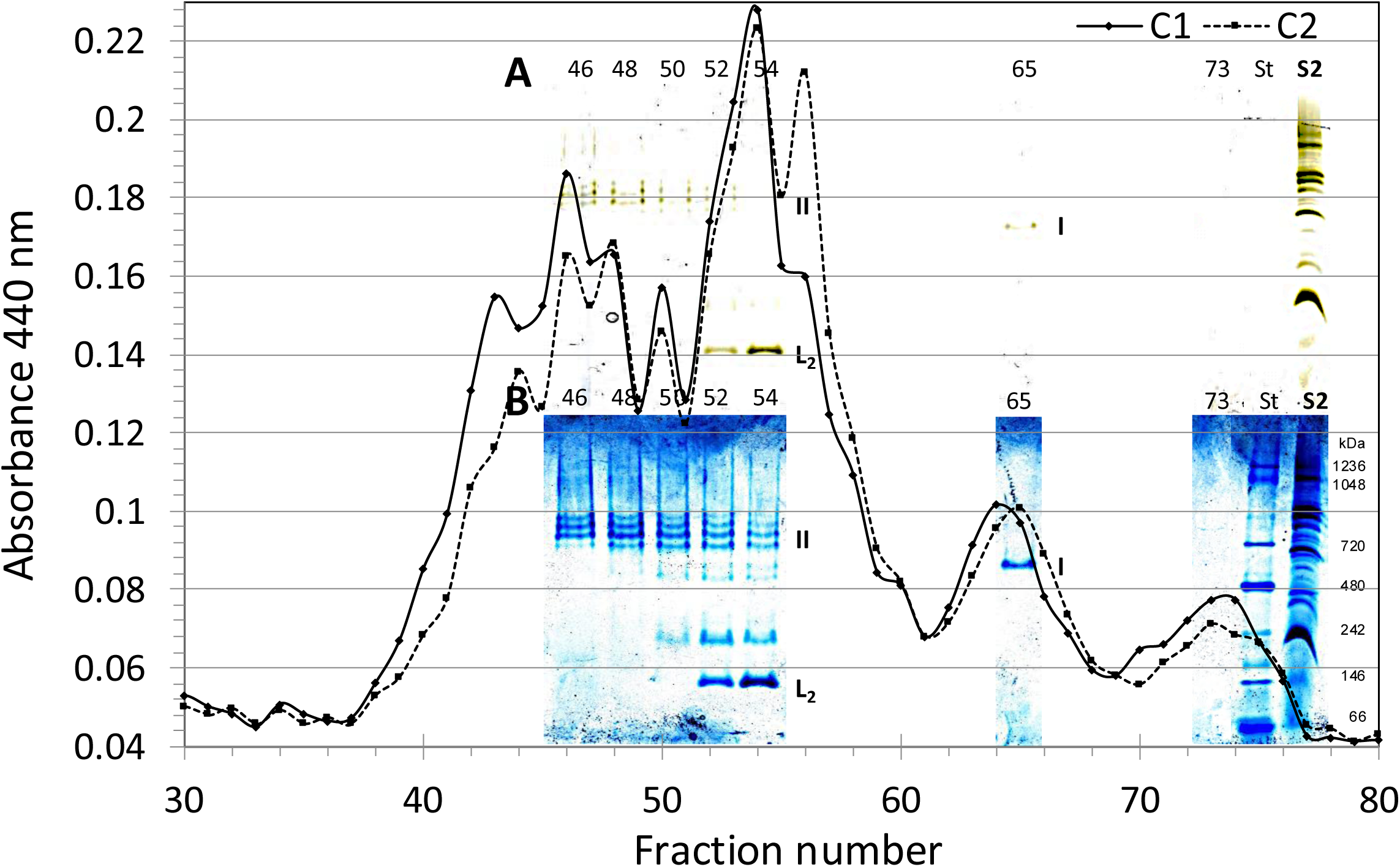
2D iZE-FFE/LN-PAGE fractionation of thylakoid membranes not extracted in step B using 20 mM β-DDM (C). The thylakoid membranes not solubilized upon extraction B (Fig. 9) were recovered by centrifugation (30 min, 25 krcf, 10 °C). Membranes were extracted using 20 mM β-DDM from the B pellet. Separation by iZE-FFE was conducted in two technical replicates, **C1** and **C2**. The two A_440_ overlay microplate scans from C1 and C2 were compared with LN-PAGE as described (Figure 8, Methods) and with unfractionated C extract **S2** and the molecular weight NativeMark standard **St**. Protein complex band labels are assigned as described (Fig. 8). The LN-PAGE analysis was conducted from fractions ∼46–54 and ∼65 plus 73.

### LN-PAGE of the three membrane protein extracts A, B and C confirms the charge order and identifies the problem of incomplete solubilization

In the three-step solubilization and one-pH step iZE, we find that both solubilization and iZE separation produce protein complex compositions consistent with experiment 6874. Changing the solubilization recipe did not remove the chemical problem: a lower detergent concentration in consecutive extracts still produces clouds of not fully solubilized protein complexes. However, separating the protein complexes by LN-PAGE improved the readability of the charge classes in the native gel, despite lowering digitonin in the first step and DDM in the second. Now, steps A and B produce a more similar overall composition than the 16 mM/8+8 mM pair in experiment 6874, but the 440 nm traces and native gels show key differences. The most important difference is in the PSI fractions. Already in the presence of digitonin only, the PSI, and the PSI–L2 band are found in extract A in the more cathodic fraction range (Fig. 7A and B, fractions 52 to 62), and the intensity of PSI is shifted with its maximum to fraction 62 (Figure 8A and B, I) instead of 55 in experiment 6874 (Figure 6A, I). In addition, lowering the digitonin concentration resolved no cytochrome *b*_6_*f* band in the PSI fractionation range but only when DDM was added in the second extraction step (Figure 8 versus 9, I, V). We note that in LN-PAGE, a higher number of PSI-specific complexes become resolved in the PSI iZE fractions (Fig. 8 and 9). In extract B, the yield of extracted PSI is increased, and here, PSI–L2 is no longer present. In the presence of DDM, PSI also shows more distinct higher-molecular-mass bands in the same FFE fractions, with a maximum in fraction 58 (Figure 9). ATP synthase is again dominant in membrane extraction A relative to B. The cloudy high-mass structures with anodic drift appear again associated with PSI in A and with PSII in B; however, in LN-PAGE the cloudy structures seen in BN-PAGE are much less prominent, while instead distinct high-molecular-mass protein complex structures become detectable in the gel stained by the Chl bound to the complexes (Figures 8-10) and which also are the basis for the absorbance traces representing the iZE graphs, after LN-PAGE. Finally, in extract C, we extract the residual membranes with 20 mM β-DDM. Here, the high detergent concentration shows that, with full solubilization, only the distinct higher-molecular-mass PSII assembly structures are observed, and the high DDM concentration is also the cause of the release of L2 bound to PSII complexes. Interestingly, not all PSII complexes in the residual grana-type membrane are equally sensitive to DDM exposure, and the iZE fractionation shows that the more anodic PSII complexes appear to bind L2 complexes more tightly. The charge separation of PSII complexes therefore indicates that the PSII complexes in the thylakoid membrane that can be solubilized by DDM are most likely structurally not alike, despite complexes showing the same molecular size in LN-PAGE (Fig. 10) and being recorded in the characteristic three-band structure of PSII resembling — C2S2M2 / C2S2M / C2S2 — structures (Caffarri et al., 2009; Järvi et al., 2011). Therefore, Cryo-TEM should be used to examine the separated charge states of photosystem complexes and assess structural differences in protein complex stability. In the DDM solubilization step, PSI can be isolated as a single peak in iZE, where the complex appears as a single band without higher-molecular-mass forms in the same iZE fraction (Figure 10).

We conclude that for each sequential extraction, the 440 nm traces of the two technical repetitions (A1/A2, B1/B2, C1/C2), and the LN-PAGE separation using LDS in the cathode buffer (Arnold et al., 2014), and the colloidal Coomassie stain of the same gel (Kang et al., 2002) show that iZE using a single pH step for protein complex separation and LN-PAGE are a superior combination of orthogonal separation techniques for analyzing the complex membrane protein composites like found in the photosynthetic membrane.

## Discussion

Native PAGE remains the right tool for asking how large a thylakoid complex is (Järvi et al., 2011; Rantala et al., 2018). It is the wrong tool for asking how charged that complex is, and it is a compromised tool for asking whether two complexes of different size and charge can survive a shared entry into polyacrylamide. iZE-FFE answers the charge question in liquid and, as a side effect, feeds the gel a simpler sample.

The scientific foundation of every separation in this paper is interval-zone electrophoresis in pH *steps*, not IEF and not continuous ZE. The acid/base pair and the detergent can be changed; the cycle cannot. HIBA/BisTris with digitonin is the Arabidopsis recipe used here (Eichacker et al., 2015). A later spinach/GDN recipe uses different chemistry on the same principle and is not included in these figures. The granted description of pH-step FFE, including the band-sharpening argument, is Weber and Weber (US 10,067,089 B2). Interval zone as a batch alternative to IEF for keeping proteins in solution was shown for a HeLa extract as P155-M (Hartmann et al., 2007). Our 2015 chapter applied that mode to thylakoids. This preprint is the charge-density, orthogonality, and solubilization-readout argument that the chapter did not make.

The BPB/DDM experiment is the most general result in the set (Figure 2). It does not depend on photosynthesis. Any membrane-protein FFE run that includes detergent is a charge-density experiment whether or not the operator plots it. Helenius and Simons (1977) used detergent-induced mobility shifts to classify proteins as amphiphilic or hydrophilic. Here the same physics applies in free solution, reminding us that the “background” of a solubilized membrane is a population of charged vesicles. Those vesicles are not invisible to electrophoresis. Digitonin does not shift BPB, and Orange G does not shift with DDM; only BPB conveyed mobility to DDM micelles. We interpret this as a partition preference, not a change in BPB charge in free solution. Digitonin has an aggregation number of about 60; its micelles have a molecular weight around 70 kDa, similar to DDM. Digitonin is structurally rigid, and the experiment indicates that different micellar packing affects BPB–micelle affinity. This is consistent with the understanding that digitonin intercalates with membrane lipids and retains annular lipids around protein complexes relative to DDM.

The comparison of SDS-PAGE (Figure 5) with native PAGE (Figure 6) of the *same* 6874 fractions is the practical result of this paper. iZE separates beautifully; that becomes apparent when the complexes are examined intact. The chemistry of sequential solubilization, however, generates an unexpected mixture of fully and only partly solubilized material. SDS-PAGE reports subunits. It therefore reports every PSI and PSII particle in a well, including the high-mass clouds, as the same polypeptide set. Native PAGE reports the particles. The high-molecular-mass PSI and PSII clouds are why the denaturing gel looks overlapped. We treat the lipid/charge interpretation of those clouds as a working model, not as a mass-spectrometric assignment. The model also accounts for a digitonin-then-DDM titration of the same membranes, in which increasing DDM collapses high-mass assemblies into discrete complex bands.

Experiment 7154 shows that lowering digitonin in step A and DDM in step B does not remove this solubilization problem, nor does operating the chlorophyll window at a single pH plateau (Figure 7). What 7154 does show is a clearer detergent series. Extract A is enriched in ATP synthase and in PSI that still binds L2, without a visible cytochrome *b*_6_*f* band in those fractions. Extract B increases PSI yield, loses the PSI–L2 band, and accumulates distinct higher-mass PSI forms; the anodic high-mass cloud tracks PSII. Extract C, 20 mM β-DDM on the residual membranes, releases LHCII from PSII, reveals the C2S2M2/C2S2M/C2S2 series, and returns PSI as a single band without higher-mass companions in the same iZE fraction (Figures 8–10). Sequential extraction is therefore not merely a way to increase yield. It is a way to ask which associations survive which detergent.

In a related experiment, we first ordered the complexes by charge in digitonin and then added extra DDM to the *same* collected wells before native PAGE. High-mass PSII–LHCII and LHCII bound to PSI fall apart; PSI cores do not (not shown). That post-iZE stripping is a different experiment from Figure 2 (BPB in empty DDM micelles) and from the HMW clouds of Figures 6–10 (incomplete solubilization *during* extraction). It matches the lipid inventory of the two photosystems: crystal structures and lipid-biosynthesis mutants show PSII as lipid-rich relative to PSI, with about 25 lipid molecules per PSII monomer versus about four per PSI (Mizusawa and Wada, 2012). Phosphatidylglycerol, DGDG, and SQDG occupy sites that stabilize the Q_B_ pocket and the binding of antenna and extrinsic subunits. DDM can compete for those sites. We therefore used LN-PAGE without adding secondary DDM, and performed solubilization steps A/B/C *before* iZE, not after.

Orthogonality of the two axes, charge and size, is the result that is specific to thylakoids. It matches the known architecture: PSII–LHCII supercomplexes of the grana are large, crowded, and, once solubilized, highly anodic under the iZE conditions used here; PSI–LHCI of the stroma lamellae is smaller in native PAGE and less anodic in iZE. ATP synthase and cytochrome *b*_6_*f* occupy intermediate or overlapping windows, which is exactly why a charge-only dimension cannot replace a size dimension — and why the combination is worth investigating. Apparent masses read from native PAGE are still apparent, as detergent, Coomassie and lipid all contribute. They are reported alongside marker positions, not as true molecular weights.

Behrens et al. (2013) reported experimental native pI values for the same Arabidopsis chloroplast complexes after digitonin solubilization, using IEF-FFE rather than iZE. Those npIs are a useful static descriptor of polypeptide charge. Our fraction numbers refer to a mobility relative to the local pH of each step, and they also include detergent and lipid associated with the photosystem particle (Helenius and Simons, 1977). In our system, the more acidic reported pI predicts a more anodic iZE window at a given pH *if* the detergent load is comparable. Behrens et al. already noted that the digitonin-to-protein ratio can shift gel mobility. Here the BPB/DDM titration makes the corresponding charge-density effect explicit in free solution. We do not convert our fraction numbers into npIs.

What this paper does not claim should be stated plainly. It does not claim that FFE of plant membranes is new: intact chloroplasts were separated in 1978. It does not claim a new photosystem supercomplex; in Yadav et al. (2017) we already took IZE fractions to the electron microscope, and the plant PSI–cytochrome *b*_6_*f* contact was described before the long-side contact later reported in *Chlamydomonas* (Steinbeck et al., 2018). It does not claim that iZE measures a physiological surface potential of the native membrane; after solubilization, the relevant object is a protein–detergent–lipid particle. It does not present mass spectrometry of the bands discussed here. Identifications are those of the published thylakoid literature cited above; the corresponding MS map of wild type versus *stn7* and *pph1* will be reported with that mutant analysis. The method does, however, show a clear separation of PSII and PSI complexes, as well as of ATP synthase and an independent pool of LHCII, and it is well placed to study regulatory changes in protein–protein interaction, for example during state transitions. Locked opposite LHCII–PSI associations are expected to change the charge axis; the method should report that.

The overlap with Eichacker et al. (2015) is intentional and limited. We reproduce Fig. 1 from that chapter as Figure 6B here. Here we report both sequential solubilizations, the 440 nm traces, the SDS versus native comparison of the same wells, and the LN-PAGE maps of a three-step extraction read for chlorophyll and after colloidal Coomassie. The findings that matter for photosynthetic membrane analysis are that a charge-first liquid dimension is orthogonal to native PAGE, that native PAGE of the iZE fractions is required to see that orthogonality, and that the product of the first dimension remains a solution. For laboratories that already run BN-PAGE, the additional hardware is a cooled FFE chamber. For laboratories that need particles for cryo-EM or native mass spectrometry, the relevant feature is that the complexes never have to be electroeluted from acrylamide.

In the presented experiments, iZE fractions were used without precipitation for (i) absorbance spectroscopy (Figures 4 and 7–10), (ii) SDS-PAGE (Figure 5), (iii) BN-PAGE (Figure 6), and (iv) LN-PAGE of sequential extracts A, B, and C (Figures 8–10). The same pipeline, applied to stroma-lamella extracts, previously yielded particles for single-particle electron microscopy of plant PSI supercomplexes, including PSI–LHCI, extra LHCII, NDH-associated PSI, and a short-side, labile contact with dimeric cytochrome *b*_6_*f* (Yadav et al., 2017). Those structures are not re-reported here. We show that iZE, especially when combined with LN-PAGE to visualize separation results, provides charge-state-based separation of complex protein mixtures, offering superior access to protein complexes for structure and function analysis. Because the protein product remains in solution, downstream processing is straightforward.

## Materials and methods

### Plant material

*Arabidopsis thaliana* was grown in soil for 3–4 weeks at 100 µmol m⁻² s⁻¹, 8 h light / 16 h dark. Rosette leaves were homogenized for 4 × 4 s in 150 mL of ice-cold 25 mM Tricine-KOH pH 7.8, 330 mM sorbitol, 1 mM Na-EDTA, 10 mM KCl, 0.15% (w/v) BSA, and 4 mM sodium ascorbate. The homogenate was filtered through Miracloth. Plastids were collected (3 min, 1 800 *rcf*), washed in 25 mM Tricine-KOH, 100 mM sorbitol, 5 mM MgCl₂, 10 mM KCl, 10 mM NaF pH 7.8 (5 min, 6 000 *rcf*), lysed on ice for 5 min in 25 mM Tricine-KOH, 5 mM MgCl₂, 10 mM KCl, 10 mM NaF pH 7.8, and membranes collected (5 min, 6 000 *rcf*). Thylakoids were suspended in TMKGS (10% (v/v) glycerol, 25 mM Tricine-NaOH pH 7.8, 100 mM sorbitol, 5 mM MgCl₂, 10 mM KCl). Chlorophyll was measured in 80% acetone (Porra et al., 1989). Aliquots corresponding to 100 µg chlorophyll were frozen in liquid N₂ and stored at −80 °C.

### Two-step solubilization, experiment #6874 (**Figures 4–6**)

Two consecutive extracts were prepared as described (Eichacker et al., 2015). Thylakoid membranes corresponding to 100 µg chlorophyll were suspended in 50 mM Tris-HCl, 5 mM MgCl₂, 10 mM KCl, 200 mM sucrose, pH 7.2, containing **16 mM digitonin**, incubated, and centrifuged. The supernatant represents solubilization 1. The membrane pellet recovered after that first centrifugation step was resuspended in the same buffer containing **8 mM digitonin plus 8 mM β-DDM**, incubated, and centrifuged; that supernatant represents solubilization 2. Samples were supplemented with 0.02% (v/v) SPADNS where noted.

### Solubilization, experiment #7154 (**Figures 7–10**)

Three consecutive extracts were produced from Arabidopsis thaliana thylakoid membranes: **A**, 10 mM digitonin; **B**, 10 mM digitonin plus 5 mM β-DDM on the A pellet; **C**, 20 mM β-DDM on the B pellet. Solubilization buffer: 25 mM Tricine pH 7.0, 250 mM mannitol, 10 mM KCl, 5 mM MgCl₂, plus the detergent of that step. Each supernatant was cleared by centrifugation (30 min, 25 krcf, 10 °C) and concentrated on 100 kDa centrifugal filters to ∼100 µl for iZE. Load each extract twice (technical replicates A1/A2, B1/B2, C1/C2).

### pI-marker mix

10 mg of the coloured pI-marker powder mix was dissolved in 1 mL NaOH, and the pH was adjusted to 8.2 (a 3–4× concentrate, diluted to 1× for the run). The standard mix contains SPADNS and amphoteric markers for pH **4.0, 4.8, 5.3, 6.4, 7.5, 8.5, and 10.1** (Šlais and Friedl, 1994). These dyes are substituted aminomethylphenols (acidic members: azo dyes; Štastná et al., 2005). They are used here to locate the pH steps (Figure 3), not to calibrate the pI of the protein complexes. Bromophenol blue (3,3′,5,5′-tetrabromophenolsulfonephthalein) was recorded by its absorbance at 595 nm in the charge-density experiment (Figure 2).

### Interval-zone FFE

Separations were run on a standard FFE system (FFE Service GmbH, Feldkirchen; BD FFE Manual 08-10-13) with a 0.2 mm gap at 10 °C. Electrode buffers: anode 100 mM H₂SO₄; cathode 100 mM NaOH, 200 mM glycine, pH ∼10. Anodic stabilization (inlet 1): 100 mM HCl, 50 mM formic acid, 50 mM HIBA, pH 3.8–4.1 (Bis-Tris). Separation buffers contained 10 mM HIBA titrated with Bis-Tris to the specific pH of 5.45, 6.21, 7.01 in the presence of 0.1% (w/v) digitonin (inlets 2-7): inlets 2–4 pH 5.4; inlets 5–6 pH 6.2; inlet 7 pH 7.0 plus 5mM NaCl; inlet 8 pH 7.0 without digitonin. Cathodic stabilization (inlet 9): 150 mM HIBA, 375 mM imidazole at pH 7.36. Counter-flow settings: 250 mM sucrose or mannitol, BisTris, AMPSO or HEPES as specified per experiment. Sample entered at inlet 7, on the cathodic side of the working pH window. Hydroxypropyl methylcellulose was omitted from the *separation media*. Both experimental settings used a 30 min 0.2% HPMC chamber coat that was washed out before buffers and sample.

The instrument was programmed for interval zone electrophoresis (Hartmann et al., 2007; Weber and Weber, 2018) at **1600 V**. Five steps: (1) **Load** — sample pump ∼70 s, media at intermediate flow, field off; (2) **Align** — sample pump off, additional transport into the field chamber; (3) **Resolve** — field on, media 40 ml h⁻¹, **4–4.5 min**; (4) **Elute** — field off, media 240 ml h⁻¹; (5) **Collect** — 96-well polyethylene plates (fraction 1 = anode). The pH steps are set in the media *before* the run and not by carrier ampholytes. For technical replicates (Figures 8–10), we loaded and separated samples consecutively. We held plates at 4 °C for native PAGE or froze them at −20 °C for SDS-PAGE.

### Experimental settings for #6874 (**Figures 4–6**)

FFE Service run sheet 6874, operator U. Sukop-Köppel. Sample: Arabidopsis WT thylakoids (S595), digitonin supernatant (plates 6874p4 / 6874p5) and residual-membrane pellet (6874p6 / 6874p7). Media as above with **0.1% (w/v) digitonin** in inlets E2–E7; **250 mM sucrose**; inlet pH 5.4 / 6.2 / 7.0. Set 1600 V / 150 mA / 250 W; displayed 1598 V / 86 mA / 137 W; 4.5 min resolution; elution field off. Chamber: 500 × 100 × 0.2 mm, 10 °C, PP60 membranes. Absorbance of 6874p4 (supernatant) was recorded at 280, 440, and 680 nm, and of 6874p6 (pellet) at 440 nm (Tecan Infinite 200). The pH of the protein plate was determined after the separation of the protein complexes, and the values are shown (Figures 4 and 7); or the pH after the pI-marker separation was analyzed (Figure 3). BN-PAGE: NativePAGE 3–12% Bis-Tris, Coomassie used in the cathode buffer, odd fractions 27–67 (gels BN4506-6874p5 and BN4507-6874p7). Panel B of Figure 6 was published as Fig. 1 as described (Eichacker et al., 2015). SDS-PAGE of the same fraction series is shown in Figure 5.

### Experimental settings for #7154 (**Figures 7–10**)

The FFE Service run sheet for experiment #7154, by the operator U. Sukop-Köppel, resembled sample S629 WT. Three-step solubilizations A, B, and C were conducted as described in the solubilization for experiment #7154; **each extract was run twice** (p6/p7 = A2, Figure 8; p8/p9 = B, Figure 9; p10/p11 = C, Figure 10). The repetitions shown in the A_440_ overlay were performed to demonstrate operative reproducibility, allowing pooling of wells. Media: 10 mM HIBA/BisTris, inlet pH 5.4 / 6.2 / 7.0, **250 mM mannitol**, **0.1% (w/v) digitonin** in E2–E7, E7 also 10 mM NaCl and used for sample inlet; upfront of buffer loading, the chamber was coated for 30 min with a 0.2% HPMC coat, then washed; counter-flow was titrated to pH 7.0. 1600 V; displayed ∼103 mA / 166 W; 4 min. pI marker on p3; pH of the sample plate is the ∼6.4 plateau in Figure 7. LN-PAGE: NativePAGE 3–12% Bis-Tris, 80 µM LDS cathode (Arnold et al., 2014); chlorophyll unstained plus colloidal Coomassie (NP4607, NP4608).

### Charge-density control (**Figure 2**)

A constant amount of bromophenol blue was dissolved in β-DDM at 0.25, 0.5, 1, 2, 4, 6, 8, and 10 mM and subjected to iZE separation, together with the pI-marker mix. Anodic displacement was read as the fraction numbers containing the BPB dye/β-DDM concentration. For parallel controls, digitonin replaced DDM, and Orange G replaced BPB.

### Concentration of iZE fractions before loading of BN- and LN-PAGE gels

180–200 µl of each fraction was concentrated to ∼25 µl using Vivaspin 100 kDa concentrators at 10 °C according to the manufacturer’s protocol.

### Native PAGE (**Figures 6**, and 8-10)

Concentrated fractions were mixed with loading dye (final 50 mM BisTris-HCl, 50 mM NaCl, 10% (w/v) glycerol, 0.001% Ponceau S, pH 7.2) and separated on Novex NativePAGE 3–12% Bis-Tris gels (Invitrogen user guide MAN0000557), 17.5 V, 10 mA, 10 W, 15–17 h. Anode and gel buffer: 15 mM BisTris, 50 mM Tricine pH 6.8 (operative pH ∼7.5; NativePAGE Bis-Tris system). The manufacturer’s cathode additive is Coomassie G-250. **BN-PAGE** (Figure 6) used 0.24 mM CBB G-250 in the cathode. **LN-PAGE** (Figures 8–10) used the same cassette, replacing the cathode additive with **80 µM (0.08 mM) LDS** (Arnold et al., 2014). Markers: NativeMark (Invitrogen) 1 236, 1 048, 720, 480, 242, 146, 66 and 20 kDa.

### SDS-PAGE (**Figure 5**)

10 µl of the FFE fraction was run on NuPAGE 4–12% Bis-Tris gels in MES-SDS (50 mM MES, 50 mM Tris-base, 0.1% SDS, 1 mM EDTA, pH 7.3) for 35–45 min. SeeBlue Pre-stained Protein Standard (3, 6, 14, 18, 28, 38, 49, 62, 98 and 188 kDa) was applied for protein mobility calibration. Gels were stained with the SilverQuest Silver Staining Kit (Invitrogen, Thermo Fisher Scientific, catalog no. LC6070) using the microwave-accelerated protocol supplied by the manufacturer.

### Staining and fluorescence

Native gels were stained with colloidal Coomassie Brilliant Blue G-250 (0.02% CBB-G250, 5% aluminum sulfate, 10% ethanol, 2% phosphoric acid) as described by Kang et al. (2002). The related high-sensitivity “blue silver” colloidal formulation is given by Candiano et al. (2004). Chlorophyll on native gels was recorded on a Typhoon Trio (excitation 633 nm, emission 655–685 nm) before Coomassie staining.

### In-gel complex labels

I, PSI (with or without LHCI); IL2, PSI binding LHCII; II, PSII (including C2S2M2/C2S2M/C2S2 and higher molecular weight multimers where resolved); IV, CF_0_CF_1_ ATP synthase holocomplex and IV1, ATP synthase head subcomplex; V, cytochrome *b*_6_*f*; L2, LHCII. M, marker; S, unfractionated sample. SDS labels in Figure 5 are assignments from subunit composition and are speculative relative to the native-PAGE assignments in Figures 6 and 8–10.

### Statistics

Technical iZE replicates (consecutive loads of the same extract) are shown where stated (Figures 8–10). The analysis did not include biological treatment statistics.

### Previously published downstream use of iZE fractionation

Single-particle EM of iZE-enriched PSI particles, reported by Yadav et al. (2017), is not part of the present dataset.

## Data availability

Figure source files and the tabulated absorbance traces will be deposited with the preprint. The 2015 protocol chapter remains the reference for the two-step digitonin recipe (Eichacker et al., 2015). The mass-spectrometry tables belong to a companion study and are not released with this preprint.

## Acknowledgments

We thank Ute Sukop-Köppel for operating the FFE and for the run sheets and Tecan plates of experiments 6874 and 7154. We thank Markart Meckel for later laboratory help at the FFE chamber. We thank Jean-David Rochaix and Geoffrey Fucile for the preparation of thylakoid membranes from *Arabidopsis thaliana* (Arnold et al., 2014). Work at UiS/IKBM was supported by the Research Council of Norway, NFR 335017.

## Author contributions

L.A.E. designed the experiments, performed the biochemical work, analyzed the data, and wrote the manuscript. G.W. developed interval-zone FFE, provided the instrument, laboratory and media, and contributed to the electrophoretic method. U. Sukop-Köppel recorded FFE operation and media for the runs shown here (experiments # 6874 and 7154).

**Supplemental Figure 1. Complete 96-well A_440_ and pH profiles after iZE of the two-step extracts (experiment 6874).** Same samples and iZE run as in Figure 4. Absorbance at 440 nm of solubilization 1 (16 mM digitonin) and solubilization 2 (8 mM digitonin plus 8 mM β-DDM) is plotted for fractions 1–96 together with the pH measured in the collection plate after the protein separation. Figure 4 shows the chlorophyll working window; this supplement shows the anodic and cathodic electrode zones.

**Supplemental Figure 2. Complete 96-well A_440_ and pH profiles after iZE of the three-step extracts (experiment 7154).** Same samples and iZE runs as Figure 7. Absorbance at 440 nm of extracts A, B, and C is plotted for fractions 1–96 together with the pH measured after the sample runs. The working window is a plateau at pH ∼6.4 (fractions 23–68); this supplement shows the full plate, including the anodic and cathodic electrode zones.

