## Supplementary figures and images for "Interval-zone free-flow electrophoresis as a charge-specific dimension for native isolation of thylakoid membrane protein complexes"

### Supplemental Figures 1 and 2

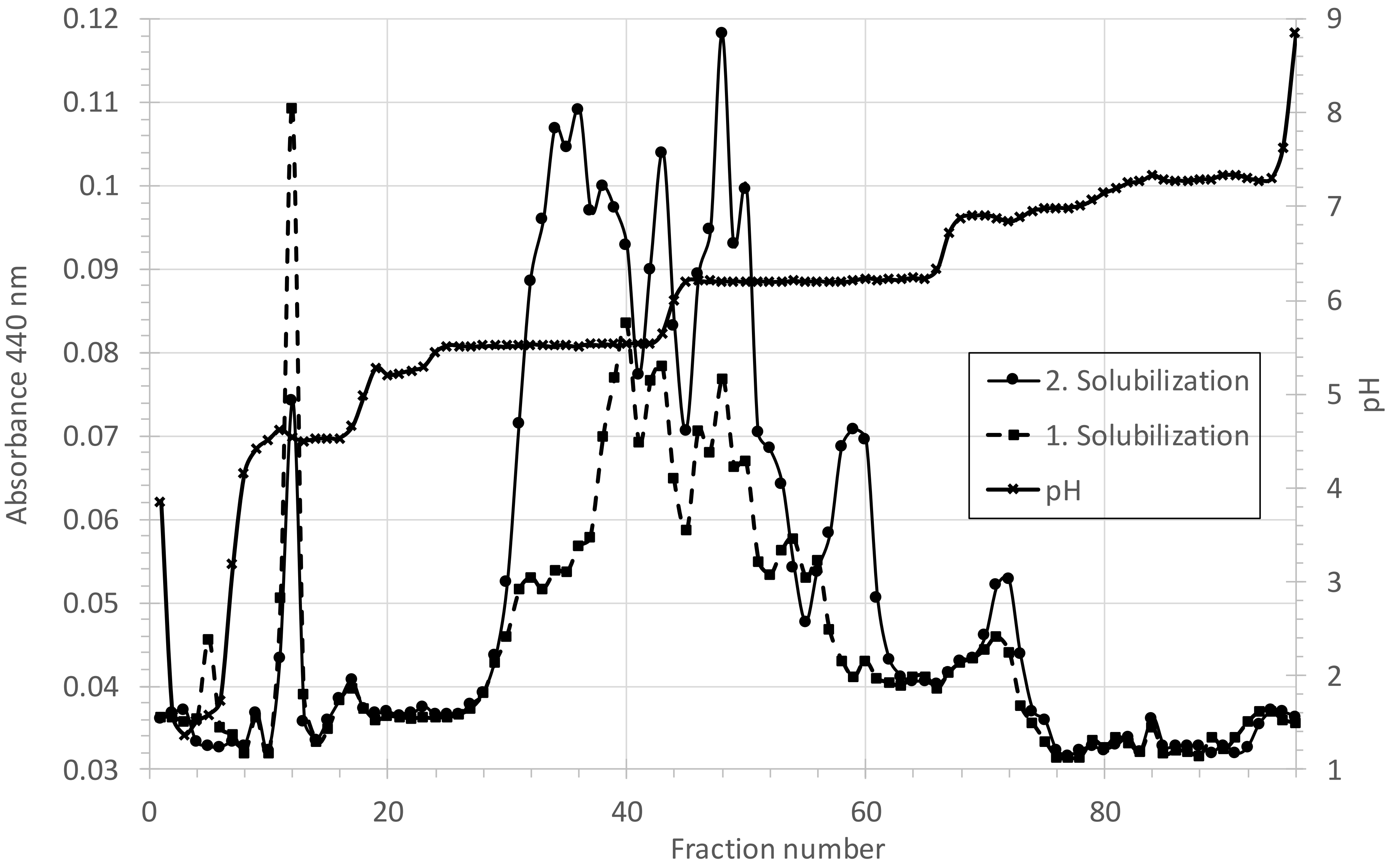

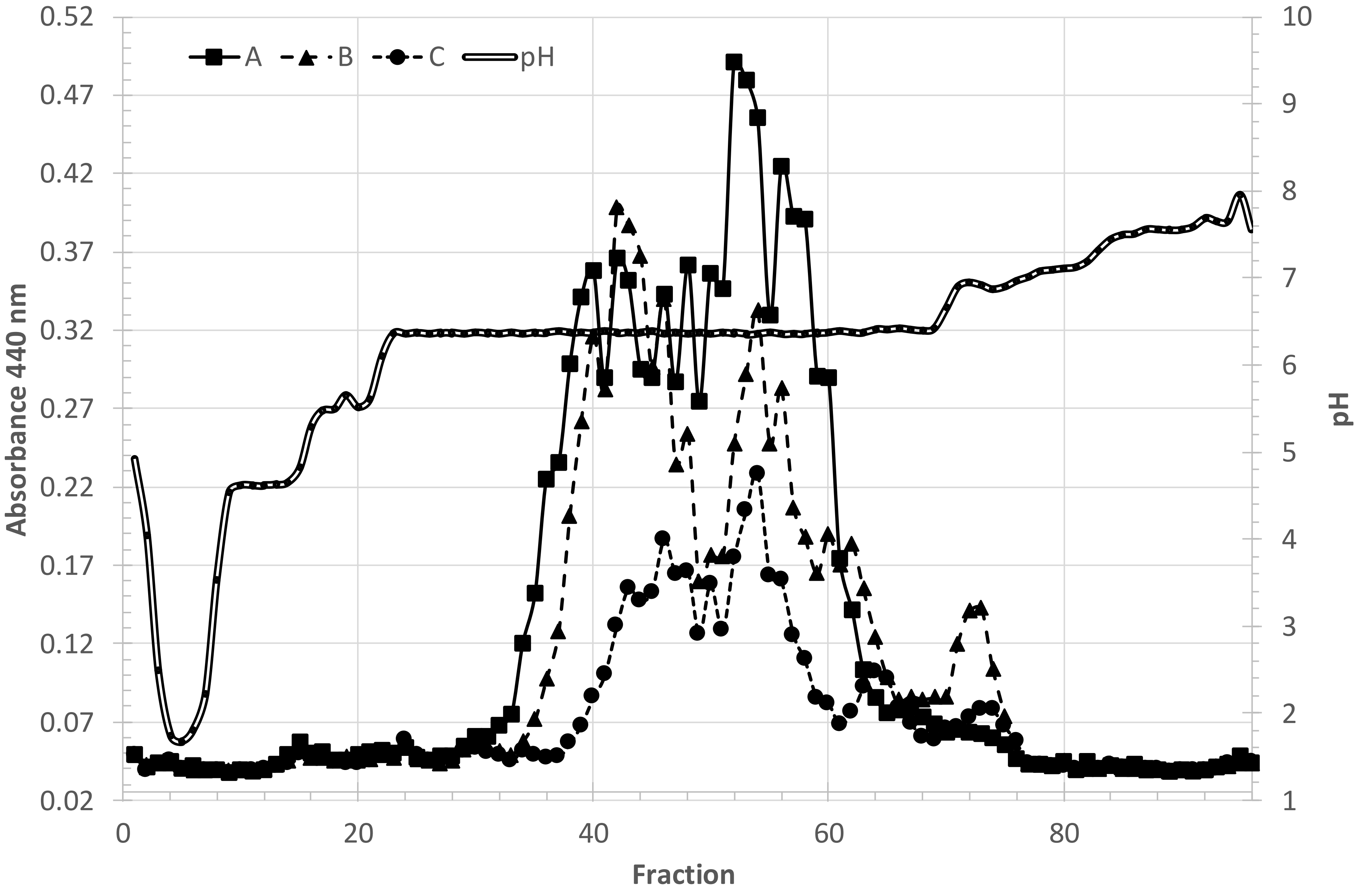
